# SARS-CoV-2 lineage assignments using phylogenetic placement/UShER are superior to pangoLEARN machine learning method

**DOI:** 10.1101/2023.05.26.542489

**Authors:** Adriano de Bernardi Schneider, Michelle Su, Angie S. Hinrichs, Jade Wang, Helly Amin, John Bell, Debra A. Wadford, Àine O’Toole, Emily Scher, Marc D. Perry, Yatish Turakhia, Nicola De Maio, Scott Hughes, Russ Corbett-Detig

## Abstract

With the rapid spread and evolution of SARS-CoV-2, the ability to monitor its transmission and distinguish among viral lineages is critical for pandemic response efforts. The most commonly used software for the lineage assignment of newly isolated SARS-CoV-2 genomes is pangolin, which offers two methods of assignment, pangoLEARN and pUShER. PangoLEARN rapidly assigns lineages using a machine learning algorithm, while pUShER performs a phylogenetic placement to identify the lineage corresponding to a newly sequenced genome. In a preliminary study, we observed that pangoLEARN (decision tree model), while substantially faster than pUShER, offered less consistency across different versions of pangolin v3. Here, we expand upon this analysis to include v3 and v4 of pangolin, which moved the default algorithm for lineage assignment from pangoLEARN in v3 to pUShER in v4, and perform a thorough analysis confirming that pUShER is not only more stable across versions but also more accurate. Our findings suggest that future lineage assignment algorithms for various pathogens should consider the value of phylogenetic placement.

## Introduction

Determining the genetic relationships between virus strains is key to SARS-CoV-2 surveillance and outbreak investigation. Lineage nomenclature systems have been a constant topic of discussion in the specialized literature with no clearly established nomenclature system for the subclassification of infectious agent lineages or subtypes and significant variability between each pathogen-specific research community (1). Early attempts at microbial lineage classification included the use of technologies such as pulsed field gel electrophoresis (PFGE) (2–6), but whole-genome sequencing (WGS) has revolutionized the field and given more discriminatory power as well as the ability to characterize additional traits such as strain antimicrobial resistance markers, virulence genes, and plasmid content in one assay (7–12). WGS has enabled phylogenetic studies to evaluate the relationships of individual sequences as a whole and allowed the development of a comprehensive lineage system classification based on genome evolution, a significant improvement over the use of single gene evolution or phenetics (13). Moreover, the use of phylogenetic tools with WGS enables improved epidemiological responses in the field by revealing viral dynamics such as in the 2013-16 Ebola epidemic in West Africa (14).

SARS-CoV-2 was first reported in December 2019, and by March 2020 it was classified as a pandemic by the World Health Organization (WHO). The rapid spread and lack of test availability in the beginning of the pandemic meant that traditional contact tracing methods were too time-intensive and inadequate to describe the scale of the pandemic. Whole genome sequencing of SARS-CoV-2 and the subsequent creation of Pango lineages (15, 16) have been central in aiding health officials to trace the spread of the virus locally and globally and identifying differences among viral lineages (17). Currently, the most commonly used tool for the lineage assignment of newly isolated SARS-CoV-2 genomes is Phylogenetic Assignment of Named Global Outbreak Lineages (pangolin), which offers parsimony-based lineage assignment using pangolin Ultrafast Sample Placement on Existing tRees (pUShER) (default on v4) and pangoLEARN (default on v3) lineage designation modes (18–20). PangoLEARN aims for a rapid assignment of lineages using a decision tree algorithm. pUShER performs a phylogenetic placement using a maximum parsimony approach to identify the lineage corresponding to a newly sequenced genome. PangoLEARN is substantially faster than pUShER. However, because the Pango lineage nomenclature system is phylogenetic (15), it is possible that pUShER is more accurate and stable in lineage assignments across subsequent releases. Given the epidemiological importance of assigning strains the correct lineages, we sought to evaluate the consistency and accuracy of the two main methods of lineage assignment currently available (21).

Despite high overall concordance between pangoLEARN and pUShER lineage assignments, pangoLEARN can be unreliable when new lineages are designated, leading to sequences that must be reassigned in a later software version despite high genome coverage/quality. In addition, greater Single Nucleotide Polymorphism (SNP) distances are found between samples assigned the same lineage by pangoLEARN (decision tree model but not random forest model). Also, more serious constellations of reoccurring phylogenetically-independent origin (scorpio) lineage call overrides are found in the pangoLEARN analyses. Therefore, we could conclude that pUShER is a more stable and an accurate method to assign pangolin lineages to SARS-CoV-2 sequences. More generally, phylogenetic placement is an appealing method for lineage assignment in rapidly evolving pathogens and should be the subject of future research for diverse pathogens.

## Materials and Methods

The genomic data used in this study were submitted to a lineage assignment pipeline using five different versions of pangolin (4 from version 3 and 1 from version 4) and MAximum Parsimonious Likelihood Estimation (MAPLE). The lineage assignments were then evaluated and compared through the methods described below (Figure 1).

**Fig. 1.**
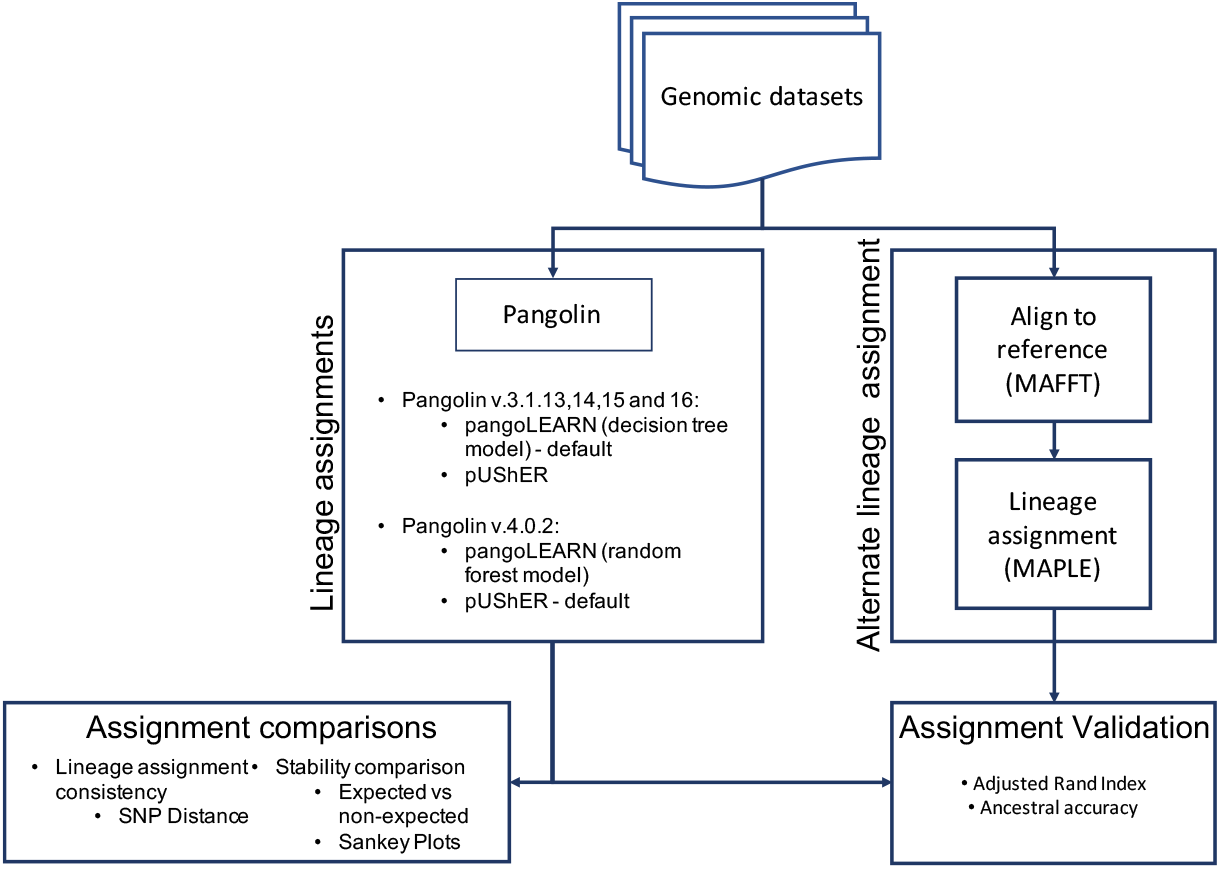
Lineage assignment and validation pipeline. Genomic datasets were run through pangolin and MAPLE for lineage assignments, assignment comparisons between pangolin assignments were performed for stability (expected vs non-expected) and lineage assignment consistency (SNP Distance), assignment validation. Assignment validation was performed comparing pangolin assignments with MAPLE assignments using Adjusted Rand Index calculations and ancestral accuracy (lineage mis-assignments belonging to ancestral or descendent lineages/sublineages).

### Data

We generated three datasets to compare the lineage assignment between pangoLEARN and pUShER as implemented in pangolin. The first (local dataset) consisted of 66,411 SARS-CoV-2 genomes collected in New York City (NYC) and the state of California (CA) with collection dates between the beginning of August 2021 and the end of November 2021: 15,862 genomes sampled in NYC by Department of Health and Mental Hygiene (DOHMH) Public Health Laboratory (PHL) and the NYC Pandemic Response Lab in addition to 50,549 genomes sampled in California via the California Department of Public Health (CDPH) COVID-Net sequencing effort. 15,393 NYC sequences and 45,326 CA sequences had a genome N percent content < 10. The second (global dataset 2021) was a random global dataset with 60,000 genomes from the National Center for Biotechnology Information (NCBI) with the same collection date range as the local dataset and sampled with equal amounts of genomes for each month with genome N percent content < 10. While random, this dataset was prone to bias due to differences in sequence deposition into NCBI impacted by factors such as total sequencing by country. The majority of genomes included came from just five countries: United States of America (31,633 sequences), England (19,097 sequences), Germany (4,677 sequences), Scotland (2,187 sequences), and Switzerland (1,662 sequences). The third (global dataset 2022) was a random global sample of 9,717 genomes from the Global Initiative on Sharing Avian Influenza Data (GISAID) collected in April 2022 with genome N percent content < 10. The composition of the third dataset skewed towards some of the same countries: United States of America (2,057 sequences), Denmark (1,730 sequences), England (1,280 sequences), Germany (1,215 sequences). However, this sampling represents a more diverse subsample than previously attained (49 versus 32 countries) and more countries with more than 100 sequences included (14 versus 7 countries), potentially due to more countries contributing sequences. This third dataset was solely used for assessing pangolin v4 given that pangolin v3 was not trained to recognize the new lineages in this dataset.

### Lineage assignments

We performed lineage assignments to the dataset sequences using five versions of pangoLEARN and pUShER. Four versions spanned Pango designation v1.2.76-93, pangolin v.3.1.13, v.3.1.14, v.3.1.15 and v.3.1.16, where pangoLEARN uses a decision tree model, and one version came from pangolin v.4.0.2/pangolin-data v1.2.133, where pangoLEARN uses a random forest model. For all the analyses, the option to assign lineages with designation hash was turned off with the option –skip-designation-hash to allow for a true comparison between both lineage designation methods.

### Lineage assignment validation

In order to validate the lineage assignments from pangolin, we used an independent method with multiple sequence alignment followed by tree search. The multiple sequence alignment was performed using Multiple Alignment using Fast Fourier Transform (MAFFT) v7.486 (22) with options – anysymbol –keeplength –6merpair –addfragments on sequences from the local dataset that had less than 10% unknown positions (N) as well as lineage consensus reference sequences (available at https://github.com/ corneliusroemer/pango-sequences) for a total of 62719 genome sequences. We implemented lineage assignment within the maximum likelihood phylogenetic software MAPLE (23) version 0.1.9 (https://github.com/ NicolaDM/MAPLE). This software was developed independently from pangolin and UShER. Placement in MAPLE was performed by inferring the joint phylogeny of reference and target genomes, and then assigning each sample to the lineage whose reference genome was the closest direct ancestor.

Disagreements between lineages assigned by MAPLE versus pangolin were scored by an in-house python script that determined whether or not one lineage was ancestral to the other, and computed the distance between the lineages as the number of edges separating the lineages on the MAPLE tree of all lineages. For example, B.1 was ancestral to B.1.2, with a distance of 1 (the edge from B.1 to B.1.2). B.1.3 and B.1.2 did not have an ancestral relationship; their common ancestor was B.1, and their distance was two (the edge from B.1 to B.1.3 and the edge from B.1 to B.1.2). We emphasize that this was not a genetic distance (*i*.*e*., the number of mutations that separate two lineages may not exactly correspond to this), but this comparison was appropriate for our analysis because we were evaluating correspondence within a lineage system. We used an in-house R script to perform an Adjusted Rand Index (ARI) comparison between the results from MAPLE and pangolin to see how consistently groups were recovered between the distinct methods.

### Pangolin versions lineage assignment comparison

We created a list of expected versus non-permitted lineage assignments between each version based on the number of newly designated lineages. We then evaluated the relationship between genome coverage and number of lineage assignments across all versions of each method and number of non-permitted lineage changes (reassignment to a non-descendant lineage in subsequent pangolin versions). We also evaluated through Sankey diagrams of lineage assignment using the five different versions of pangolin (v3.1.13.,v3.1.14,v3.1.15,v3.1.16 and v4.0.2) to look at the distinct pattern of assignment and reassignment (including non-permitted lineage changes) within pUShER and pangolin versions.

### SNP Distance between samples

A pairwise SNP distance matrix was created using snp-dists v.0.8.2 (https://github.com/tseemann/snp-dists). For each version of pangolin tested and each dataset, we calculated the SNP distances between all the sequences that were designated a lineage name.

## Data availability

The data used in this manuscript is publicly available at GISAID and NCBI. Sequence data with less than 90% genome coverage that were not found in public databases were made available in a public github repository. The list of the accessions along with metadata and genomic sequences of the global and local datasets used in this study are available on github at url https://github.com/nychealth/COVIDconsensus-genomes-pangolin-analysis.

## Results

### Dataset sampling

To evaluate the performance of pangolin calling by either pangoLEARN or pUShER, we built two initial datasets: a local US dataset that consists of 60,719 samples from California and New York City and a 60,000 sample global dataset. The timeframe of sampling was August to November 2021 to avoid early sequencing quality concerns surrounding the newly designated Omicron strains. Therefore, samples included in this study were largely Delta sublineages (Figure 2). The US dataset was more deeply sampled and could potentially magnify lineage assignment errors that apply only to a small set of samples, thus the performance against a global dataset was also necessary. It is worth noting that the majority of the global dataset consisted of sequences from the US and England and was a biased subsampling of globally circulating strains.

**Fig. 2.**
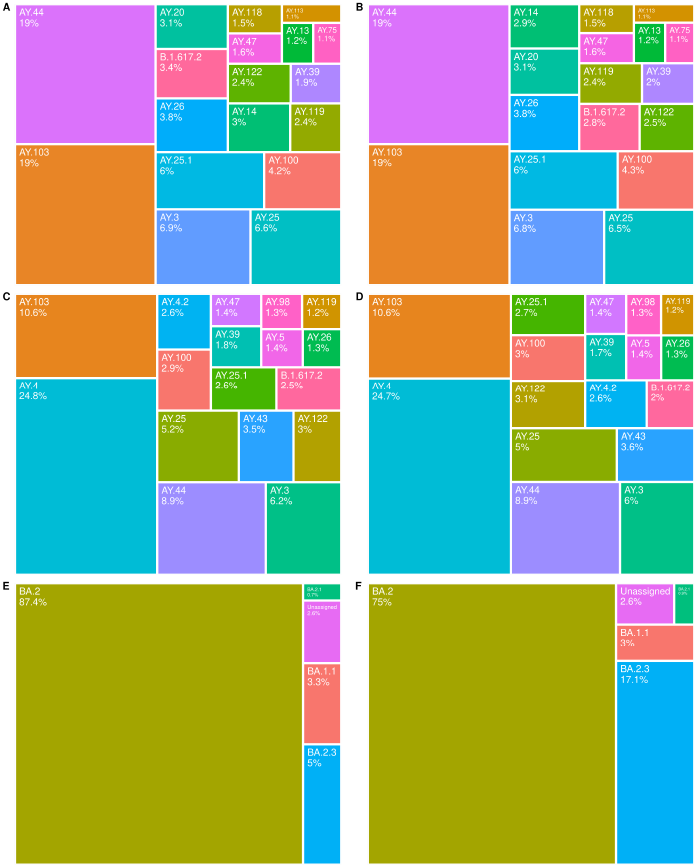
Lineage calls of datasets as determined by pUShER (A,C,E) and pangoLEARN (random forest model) (B,D,F) in pangolin v.4.0.2. A/B: local dataset, C/D: 2021 global dataset, E/F: 2022 global dataset. For A-D, only lineages present at 1% prevalence in the dataset are shown and for E-F, .01%.

There are differences in sublineage prevalence between the local and global datasets that reflect the effect of location on circulating variants. While countries can also enter different stages of the SARS-CoV-2 pandemic at different times, e.g. B.1.1.7 emerged in the UK before being found in other countries, the time period studied began in the middle of Delta’s dominance (roughly May/June-December 2021) and was not a significant contributing factor.

After the release of pangolin v4, we created a third global dataset of 9,717 sequences from April 2022 to evaluate performance of a newer methodology employed by pan-goLEARN. This subset was still biased towards certain countries like the US, England, Germany, and Denmark and reflected the waning of BA.1, which dominated during the early Omicron wave, and the subsequent takeover of BA.2. Additionally, it was a less diverse subset of sublineages than the previous datasets and performance on this subset may not have necessarily been generalizable.

### Overall Concordance and SNP Distance

The concordance of the methods improved with each new model (Table 1). This was more apparent with the 2021 global dataset and was likely due to AY.4 prevalence. Overall, the two methods were highly concordant and differed mostly on sublineage calls.

**Table 1.** Agreement of calls of pangoLEARN and pUShER across the local and global datasets for different Pangolin versions (v3.1.13, v3.1.14, v3.1.16 and v4.0.2).

|  | Datasets |  |  |
| --- | --- | --- | --- |
| Pangolin version | Local | Global 2021 | Global 2022 |
| v3.1.13 (decision tree) | 84.68% | 82.13% | N/A |
| v3.1.14 (decision tree) | 87.85% | 84.50% | N/A |
| v3.1.15 (decision tree) | 91.02% | 90.55% | N/A |
| v3.1.16 (decision tree) | 92.93% | 89.45% | N/A |
| v4.0.2 (random forest) | 97.28% | 97.35% | 86.9% |

Pangolin v4 released in April 2022 changed the pangoLEARN model to a random forest, fixed the previous calling errors of pangoLEARN (decision tree model) and was not significantly different from pUShER in any of the historical datasets. This was to be expected as similar sequences from those datasets were likely included in the training datasets for the new model. pangoLEARN (random forest model) v4 had good concordance with pUShER on a contemporary dataset collected in April 2022 but it was much lower than with the historical datasets. The lower concordance could be largely attributed to pUShER assigning BA.2 and pangoLEARN BA.2.3. Assigning lineages with a newer version of pangoLEARN (random forest model) v.4.1.2 (UShER-v1.13 and pangoLEARN-v1.13) resolved the assignment in the favor of pUShER as pangoLEARN BA.2.3 calls decreased significantly from 1,662 to 392, while pUShER BA.2.3 calls changed minimally from 483 to 395. Thus, the new pangoLEARN (random forest model) method may still be prone to certain types of miscalling.

Because pangolin lineages are defined phylogenetically, we expected sequences that were given the same lineage designation to be more closely related than those of a different lineage. Pairwise SNP distances were calculated, and the average pairwise distance per lineage is shown in Figure 3. It is evident that as the Delta wave progressed and more sublineages evolved, pUShER was able to call these newly defined sublineages with a lower average SNP distance between samples compared to pangoLEARN (decision tree model). This was true for both the local and global 2021 dataset with the difference between the two methods larger within the global. This was likely due to the high prevalence of AY.4 in the global 2021 dataset, which was known at a previous date to be overcalled by pangoLEARN (see https://github.com/cov-lineages/pango-designation/issues/221). In general, pangoLEARN (decision tree model) showed improvement over the course of the Delta wave by shrinking the average SNP distance between lineages with each successive model. Notably, pangoLEARN v4 (random forest model) did not differ significantly from pUShER in any of the datasets.

**Fig. 3.**
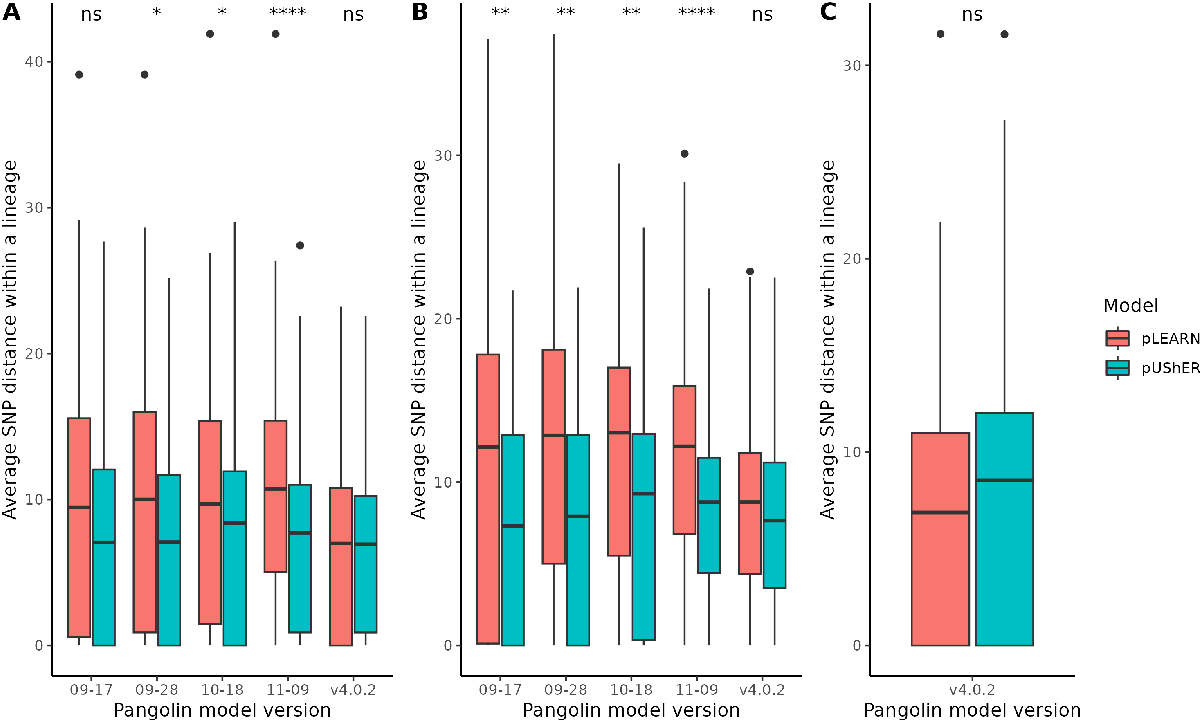
Average SNP distance between pangoLEARN and pUShER on pangolin. A: local dataset, B: 2021 global dataset C: 2022 global dataset. ns = not statistically significant / * = 0.05 / ** = 0.01 / **** = 0.0001. 09-17, 09-28, 10-18, and 11-09 used the pangoLEARN decision tree model, and v4.0.2 used the pangoLEARN random forest model.

### Scorpio Analysis

Scorpio takes a set of lineage-defining “constellations” with rules to classify each sequence by its specific mutations. It is manually curated and is limited to the WHO Variants of Concern, Variants of Interest, or Variants Under Monitoring (24). Prior to pangolin v4.1, when pangoLEARN or pUShER made an assignment that conflicted with scorpio’s assignment, pangolin overrode the pangoLEARN or pUShER assignment with the scorpio assignment. This allowed pangolin to make higher accuracy assignments when new lineages emerged (pangoLEARN would initially lack sufficient training data for new lineages). However, this proved problematic with the emergence of BA.4 and BA.5 which saw scorpio overriding correct assignments of these lineages and outputting BA.2 as the lineage (see https://github.com/cov-lineages/scorpio/issues/47). This scorpio issue was due to early BA.4 and BA.5 having lower quality and lacking lineage defining mutations. However, due to the timeframe sampled, BA.4 and BA.5 sequences were not present in our 2022 global dataset. No other large scale problems with scorpio have previously been documented. Therefore, scorpio overrides in this study can be evaluated as erroneous calls made by pangoLEARN or pUShER.

In general, scorpio was not a significant contributor to the overall accuracy of these methods as it was usually used in less than 1% of cases. Comparing the two methods, scorpio overrode pangoLEARN calls more often than pUShER calls (Figure 4). As expected, scorpio was used to correct many incorrect AY.4 calls and accounted for the largest difference between the two methods (pangolin model 10-18). We observed pangoLEARN (decision tree model) erroneously calling B.1/B.1.1 sequences when pUShER did not have a similar problem. Given the timeframe of the specimens (Delta or Omicron wave), it was reasonable to not expect these sequences to be present, and this was corroborated by the genomic mismatches flagged by scorpio in these sequences. B.1 miscalls were mostly present in pangolin model 9-17 and to some extent model 09-28. This reappeared as an issue in pangoLEARN (random forest model) v4 with B.1.1.

**Fig. 4.**
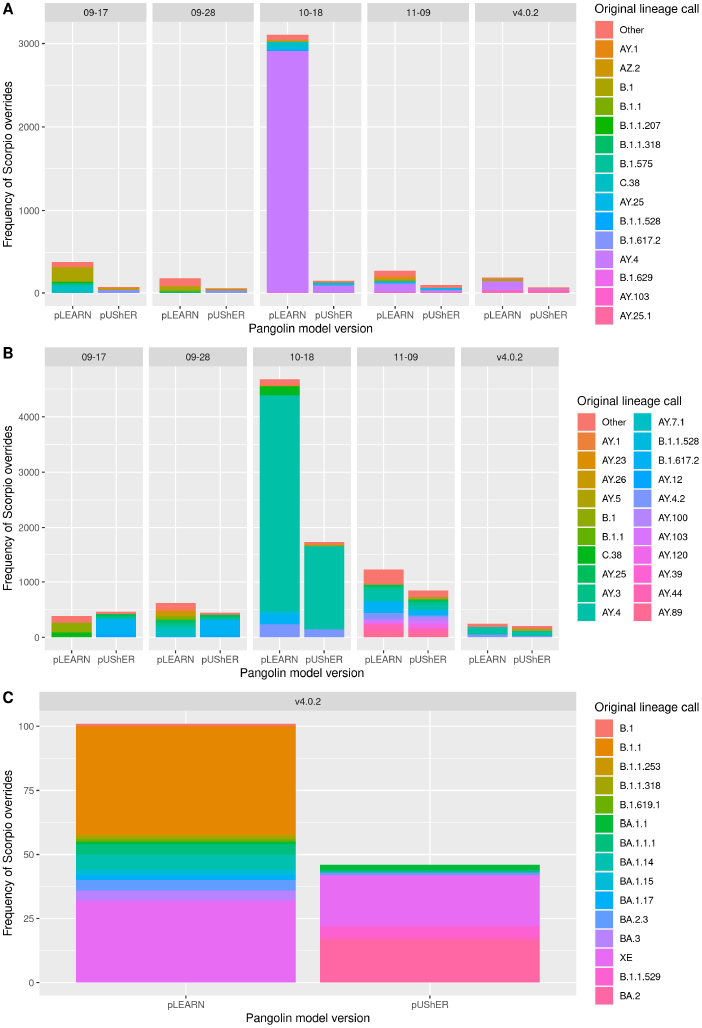
Number of scorpio overrides between pangoLEARN and pUShER on Pangolin. A: local dataset B: 2021 global dataset, C: 2022 global dataset. 09-17, 09-28, 10-18, and 11-09 used the pangoLEARN decision tree model, and v4.0.2 used the pangoLEARN random forest model.

Initially, lineage categorization was performed using phylogenetic tree search methods such as IQTree. However, as the number of sequences grew, these methods became unfeasible. To address this issue, UShER became the go-to method for tree search and lineage categorization. This shift may explain why pUShER showed a higher degree of accuracy in lineage assignments compared to pangoLEARN.

### Allowed reassignments

The Pango lineage system was explicitly designed to be updated with the SARS-CoV-2 pandemic as the virus continues to evolve (15). For that reason, some lineage reassignments were expected for a given genome sequence. For example, a sample originally designated in one lineage might be reassigned to a new daughter lineage of its original assignment. This occurs as an expected part of the Pango system when new lineages are designated. However, in some cases, a sample may be reassigned to a non-descendant lineage in subsequent versions of pangoLEARN or pUShER due to error in the assignment approach. We will refer to such lineage reassignments as non-permitted lineage changes. Instability in lineage assignments might cause problems for interpretation that relies on precise and reliable lineage definitions for individual samples.

The consistency of assignments by pangoLEARN was inferior to pUShER. Even though 81% of the sequences being assigned by pangoLEARN had a maximum of two calls across different pangolin versions, pUShER outperformed pangoLEARN by assigning 97% of the sequences a maximum of two calls (Table 2). Although a large number of calls across different versions of pangoLEARN could be associated with the designation of new lineages, 27% of the sequences analyzed with pangoLEARN presented at least one non-permitted change, while only 7% of sequences assigned by pUShER present at least one non-permitted change (Table 3 and Figures 5 and 6). While pangoLEARN v4 (random forest model) was included in this analysis, the results were a reflection of the instability of pangoLEARN v3 (decision tree model) and cannot be generalized to v4.

**Table 2.** Number of lineage assignments using pangoLEARN and pUShER across five different versions (v3.13,v3.v3.14,v3.15,v3.16 and v4).

| application\n of calls | 1 | 2 | 3 | 4 | 5 |
| --- | --- | --- | --- | --- | --- |
| pangoLEARN | 19222 (29%) | 34463 (52%) | 9537 (14%) | 2515 (4%) | 673 (1%) |
| pUSHER | 21444 (32%) | 43229 (65%) | 1619 (2%) | 96 (>0%) | 22 (>0%) |

**Table 3.** Number of non-permitted lineage changes using pangoLEARN and pUShER across five different versions (v3.13,v3.v3.14,v3.15,v3.16 and v4).

| application\non-permitted changes | 0 | 1 | 2 | 3 | 4 |
| --- | --- | --- | --- | --- | --- |
| pangoLEARN | 48586 (73%) | 10706 (16%) | 4170 (6%) | 2072 (3%) | 876 (1%) |
| pUSHER | 61518 (93%) | 3964 (6%) | 670 (1%) | 197 (>0%) | 61 (>0%) |

**Fig. 5.**
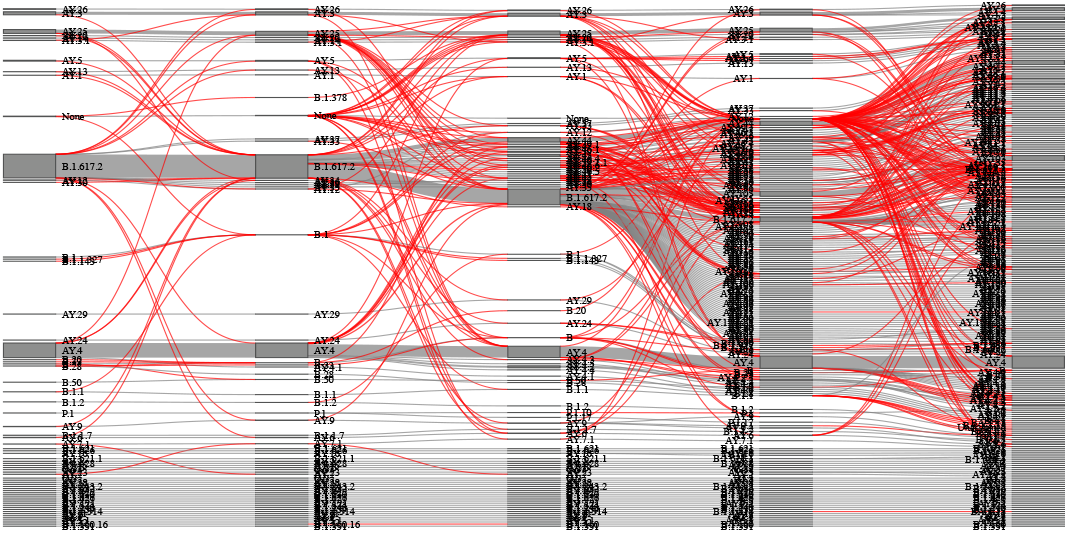
Sankey diagram of lineage assignment using pUShER across five different versions. Each column represents one version of pangolin in order of release (v3.1.13.,v3.1.14,v3.1.15,v3.1.16 and v4.0.2). Red lines represent unexpected changes between versions.

**Fig. 6.**
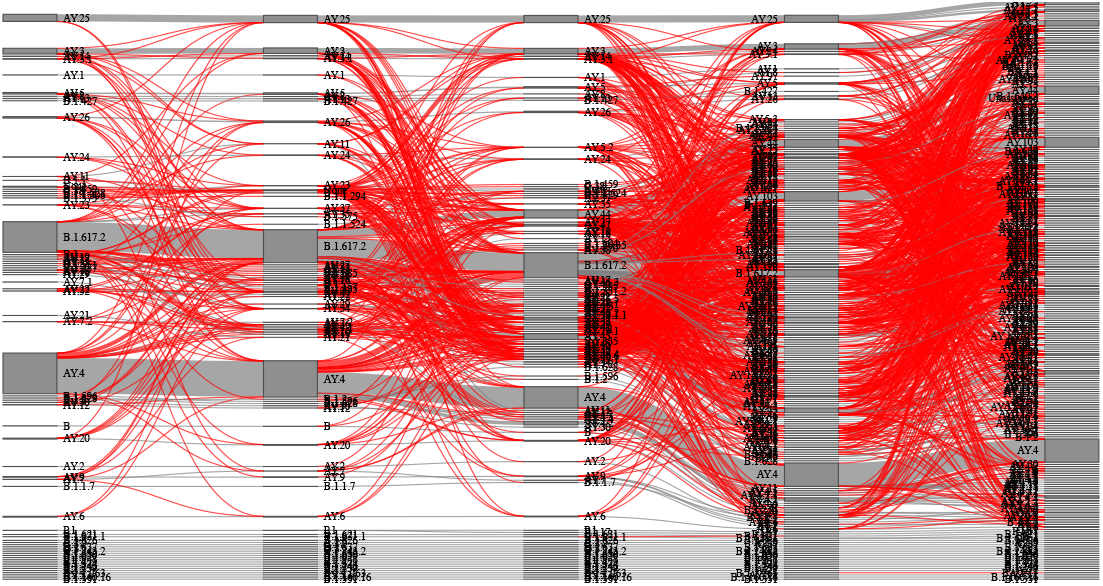
Sankey diagram of lineage assignment using pangoLEARN across five different versions. Each column represents one version of pangolin in order of release (v3.1.13.,v3.1.14,v3.1.15,v3.1.16 and v4.0.2). Red lines represent unexpected changes between versions.

Furthermore, the number of pangoLEARN lineage assignments for a given sequence seemed to be independent of the genome coverage whereas pUShER assignments had a higher number of non-permitted lineage changes and consequently higher number of lineage assignments as the genome coverage decreased (Figure 7). This reflects the expected higher phylogenetic placement uncertainty of less complete genomes.

**Fig. 7.**
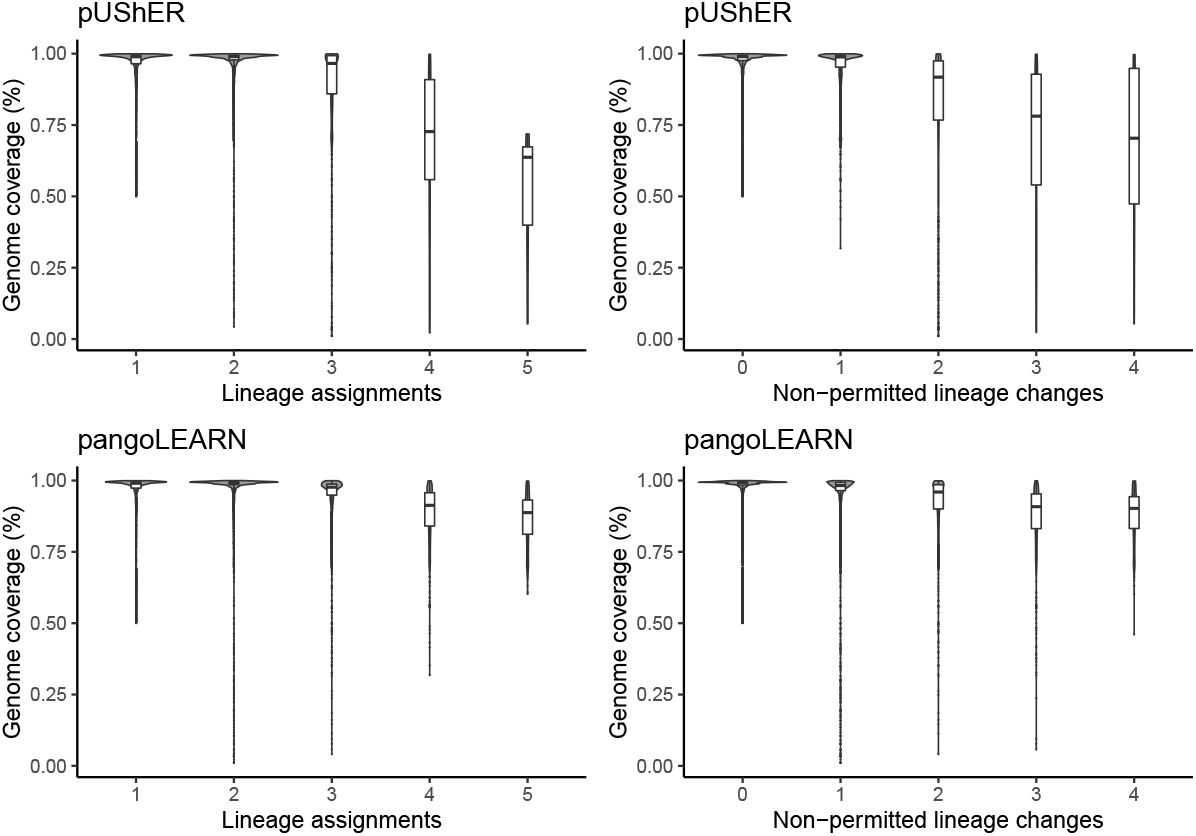
Violin plot of sample distribution based on genome coverage and pangoLEARN/pUShER lineage assignments for all CA and NYC samples. A: pUShER lineage assignment calls and reference genome coverage; B: pUShER lineage assignment non-permitted lineage changes and reference genome coverage; C: pangoLEARN lineage assignment calls and reference genome coverage; D: pangoLEARN lineage assignment non-permitted lineage changes and reference genome coverage.

### Lineage assignment validation

As the pango lineage designation system is phylogenetic in nature, we wanted to benchmark the results of v4 pUShER and v4 pangoLEARN (random forest model) against a full maximum-likelihood phylogenetic method, MAPLE (23). The MAPLE lineage assignment recovered 91.44% of the calls made by pUShER and 89.56% of the calls made by pangoLEARN (Table 4). The calculated ARI for both comparisons was above 0.90 indicating excellent recovery of similar clusters of lineage assignments regardless of the specific lineage call made for each sequence.

**Table 4.** Validation of pangolin v.4 lineage assignments (pUShER and pangoLEARN) by comparison with MAPLE lineage assignments. Match and mismatch = Direct comparison of matches/mismatches of main call between the distinct methods. ARI = Adjusted Rand Index (>0.90 = excellent recovery).

|  | **MAPLE vs pUShER** | **MAPLE vs pangoLEARN** |
| --- | --- | --- |
| Match | 55522 (91.44%) | 54379 (89.56%) |
| Mismatch | 5197 (8.56%) | 6340 (10.44%) |
| ARI | 0.935827881 | 0.925234854 |

When looking into the mismatches, we found that 5537 (87.3% of mismatches) of MAPLE vs pangoLEARN (random forest model) mismatches were ancestrally related calls with distances of 1 or 2 sublineages between the calls, while 803 (12.7% of mismatches) were siblings with distances of 2 to 4 to their common ancestor, and the presence of a mismatch due to a recombinant (XB) (Supplementary table 1). For MAPLE vs pUShER, a similar ratio was found, with 4,516 (86.9%) being ancestrally related with distance of 1 or 2 sublineages between the calls and 681 (13.1%) being siblings with distance of 2 to 4 from their common ancestor in addition to the presence of a mismatch due to a recombinant (XB) (Supplementary table 2).

We looked at the unique individual pairs of mismatches between pangolin methods and MAPLE. Of 331 pairs of disagreeing lineage assignments, 173 had an ancestral relationship while 158 did not. 144 pairs had a distance of 1 node in the tree to each other, 144 had a distance of 2, 25 had a distance of 3, 16 had a distance of 4 and, in two cases, one lineage but not the other was a recombinant (XB), indicating that half of the unique pairs of mismatches were close calls.

We also included the alternative assignments for both pUShER and pangoLEARN (random forest model) of those sequences that had mismatches with MAPLE (Supplementary table 3). Our main finding was that the presence of one disagreeing non-ancestral pair, AY.25.1 from MAPLE vs. AY.4 from pangolin, was caused by pangolin v4.0.2’s use of scorpio to override pUShER and pangoLEARN (random forest model). 20 sequences assigned AY.25.1 by MAPLE were also assigned AY.25.1 by pUShER and pangoLEARN, but were assigned AY.4 by scorpio, which overrode the pUShER and pangoLEARN assignment in the pangolin output. Together, these results show that there is high concordance between pangolin methodology and an independent phylogenetic-based method, giving confidence that pangolin can account for the phylogenetic structure underlying SARSCoV-2 evolution.

## Conclusions

Given the increased stability and reduced rate of nonpermitted lineage reassignments by pUShER coupled with its higher reliability when analyzing high genome coverage/quality sequences compared to pangoLEARN v3 (decision tree model), we recommend that the pUShER option be selected as the first choice when assigning lineages to newly and previously sequenced genomes. While we hypothesize pangoLEARN v4 (random forest model) is more stable than v3, UShER has still been shown to have fewer scorpio overrides and is likely to be as or more stable. We stress, however, there are two important caveats to this recommendation.

First, the lineage system is explicitly phylogenetic and recently the Pango curation team has used a tree inferred with pUShER to assign new lineages. Thus, if there are systematic biases associated with phylogenetic inference in pUShER, then consistently inaccurate phylogenetic placements might create spurious lineage designations and assignments that appear to be consistent lineage calls. We consider this unlikely because the pUShER’s accuracy for SARS-CoV-2 phylogenetic inference was quite high (18, 25).

Second, lineage assignment with pUShER had higher computational cost than with pangoLEARN; however, the compute costs associated with lineage assignment were relatively small compared to the total costs associated with producing a single genome sequence. Furthermore, pUShER could efficiently exploit parallelism to decrease runtime. We believe that the advantages of increased stability of lineage assignments outweigh marginal additional compute costs except possibly when reanalyzing vast datasets on a regular basis as is done with large repositories such as GISAID. However, a complete reanalysis of 6 million genomes would cost approximately $43.44 on a typical cloud instance (see https://github.com/cov-lineages/pangoLEARN/issues/32#issuecomment-946937425) if efficiently exploiting multi-core architectures. This suggests that cost is still a relatively minor consideration when choosing lineage assignment modes.

As of pangolin v4, pUShER is now the default option due to its accuracy and performance, which has been verified by benchmarking against the tree created by MAPLE. Changes have also been made to potentially decrease runtime through the implementation of an assignment cache (–addassignment-cache and –use-assignment-cache). In addition, as of v4.1, scorpio no longer overrides pUShER lineage assignments but continues to do so for pangoLEARN. Outbreak investigations are a case study in how pUShER’s high accuracy and robustness can reduce superfluous resource consumption. Lineages called by pUShER can be trusted to cluster together on a phylogenetic tree and thus be genetically similar, avoiding chasing unrelated cases and maximizing resources allocated to contact tracing. Similarly, rapidly increasing sublineages can be scrutinized for higher fitness and/or immune evasion if the lineage calls can be trusted to be reliable. Thus, for newly emerging pathogens undergoing rapid evolution, our results suggest that phylogenetic placement is a superior option for lineage assignment than machine learning methods.

## Acknowledgements

We would like to acknowledge Rachel Colquhoun (University of Edinburgh) who has done most of the work on Scorpio/constellations as used by pangolin; the NYC DOHMH PHL staff; the CDPH California COVIDNet Team and these CDPH members: Dr. Kathleen Jacobson, Dr. Carol Glaser, Dr. Mayuri Panditrao, Dr. Christina Morales, Dr. Nikki Baumrind, Elizabeth Baylis, Sabrina Gilliam; the University of California Office of the President, and COVIDNet WGS Lab Partners throughout California. COVID sequencing in NYC was supported (in part) by the Epidemiology and Laboratory Capacity (ELC) for Infectious Diseases Cooperative Agreement (Grant Number: ELC DETECT (6NU50CK000517-01-07) funded by the Centers for Disease Control and Prevention (CDC). The manuscript contents are solely the responsibility of the authors and do not necessarily represent the official views of CDC or the Department of Health and Human Services. Genomic surveillance by COVIDNet was funded by Centers for Disease Control and Prevention, Epidemiology and Laboratory Capacity for Infectious Diseases, Cooperative Agreement Number 5 NU50CK000539. The findings and conclusions in this article are those of the authors and do not necessarily represent the views or opinions of the California Department of Public Health of the California Health and Human Services Agency.

